# Mathematical modeling and simulations for large-strain J-shaped diagrams of soft biological tissues

**DOI:** 10.1101/275206

**Authors:** K. Mitsuhashi, S. Ghosh, H. Koibuchi

## Abstract

Herein, we study stress-strain diagrams of soft biological tissues such as animal skin, muscles and arteries by Finsler geometry (FG) modeling. The stress-strain diagram of these biological materials is always J-shaped and is composed of toe, heel, linear and failure regions. In the toe region, the stress is zero, and the length of this zero-stress region becomes very large (≃ 150%) in, for example, certain arteries. In this paper, we study long-toe diagrams using two-dimensional (2D) and 3D FG modeling techniques and Monte Carlo (MC) simulations. We find that except for the failure region, large-strain J-shaped diagrams are successfully reproduced by the FG models. This implies that the complex J-shaped curves originate from the interaction between the directional and positional degrees of freedom of polymeric molecules, as implemented in the FG model.

## I. INTRODUCTION

Biological materials such as muscles, tendons and skin are known to be very flexible and strong, and for this reason, these materials have attracted considerable interest with regard to the design of artificial materials or meta-materials [1, 2]. The mechanical properties of these materials are of fundamental importance in their applications. The stress-strain diagram is a typical approach for characterizing the mechanical strength of these materials, and numerous experimental studies on this topic have been conducted [3–17].

It has been experimentally observed that the stress-strain diagram of soft biological tissues is J-shaped and that the curve is composed of toe, heel, linear and rapture (or failure) regions [3–17]. The failure region is beyond the scope of this paper and will not be taken into consideration. In the toe region, the stress is almost zero, implying that the materials freely extend without external forces, similar to the behavior of the soft-elasticity region of liquid-crystal elastomers [18–24]. The existence of this zero-stress region is the reason why we call the curve Jshaped. The origin of the shape of curve is qualitatively understood such that a variety of materials, e.g., myosin, titin, and collagen, and structures, e.g., the cross-linked, entangled or network structures of these materials, contribute to this phenomenon, and hence, the interaction range extends to different length scales similar to a multicomponent material [25]. The continuum mechanics approach based on the strain energy functional also successfully describes the J-shaped diagrams [5]. In the context of continuum mechanics, the non-linearity in the J-shaped curve is understood as hyperelasticity [26, 27]. However, the mechanism of the interaction underlying the J-shaped curve is still unclear on a quantitative and a microscopic level.

In Refs. [28, 29], we studied J-shaped curves by the Finsler geometry (FG) modeling technique and obtained Monte Carlo (MC) data consistent with previously reported experimental results, in which the toe length is up to 40% ∼ 50% on the strain axis. For these experimental J-shaped curves of small toe length, FG modeling successfully describes the diagrams.

However, the toe length reaches 150% in some biological materials [7]. The length of the toe region is generally, albeit not always, limited to less than 50% [3–7]. One of the reasons for such a large difference in the toe length comes from the difference in the component molecules. The main component of materials with a small (large) toe length is the collagen (elastin, etc.) molecule [1]. Moreover, the hierarchical structure, such as that in the collagen molecule, is also expected to be one of the reasons for the variety of toe lengths [2]. Due to this hierarchical structure of molecules, there must be a considerable difference in the interactions between the hierarchies. The mechanism for large toe-length materials is qualitatively different from that of small toe-length ones. As a result, we should check whether the J-shaped diagrams with a large toe length can be studied by the FG modeling technique or not.

In this paper, the existing experimental J-shape curves of biological tissues such as animal skin, muscles and arteries are compared with the simulation results. The experimental curves of these materials are grouped into two types: the group of diagrams with a small heel (Sheel) and the group of diagrams with a large heel (L-heel) (Figs. 1(a),(b)) [4–7]. The diagram with an S-heel (Fig. 1(a)) is decomposed into two straight lines with different slopes, while the diagram with an L-heel is decomposed into two different linear lines and a curve for the heel between the two lines. Moreover, close to the failure region, some of the curves have a convex part where the collagen fibers start to break (Fig. 1(b)).

**FIG. 1.**
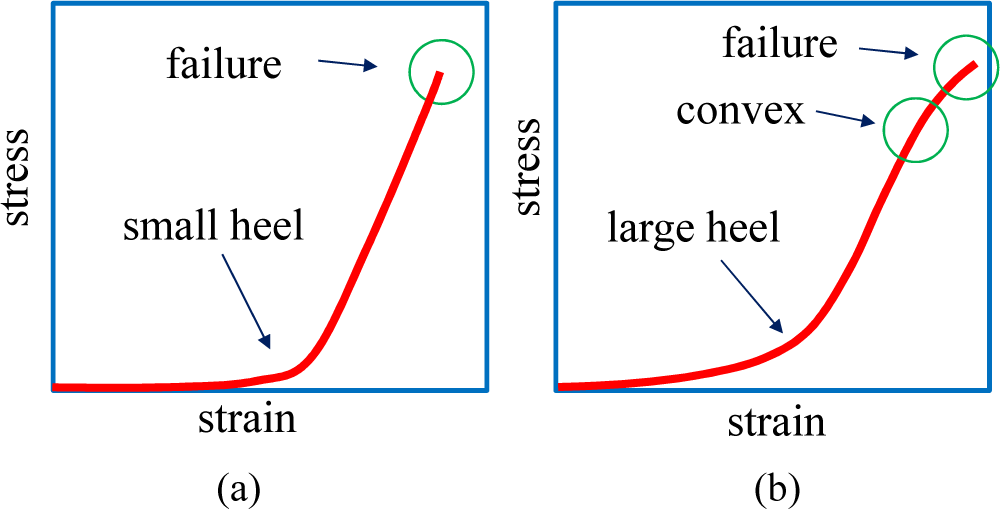
A J-shaped diagram is composed of two different linear lines, and the region where two lines are smoothly connected is called the heel. The J-shaped diagrams are decomposed into two groups: the diagrams with (a) a small heel and (b) a large heel. The curve in (b) is convex upward in the large-strain region and hence is highly non-linear.

We show that the curve with an S-heel is successfully reproduced by our two-dimensional (2D) FG model, while the curve with L-heel is not described by the 2D FG model but can be reproduced by a 3D FG model. The large strain curves with a convex part are found to be well-fitted by 2D FG model data. Our results in this paper indicate that the FG modeling technique can be used to analyze a wide range of J-shaped stress-strain diagram of biological tissues such as tendon, skin, muscles and arteries. Since the main component of these materials is a polymer such as collagen fiber, the polymeric degrees of freedom can be coarse grained and are simply replaced by the variable *σ*(*∈ S*^2^: unit sphere) in the FG model. This simple coarse graining is key to understanding the mechanical properties of these biological materials mathematically.

We comment on the reason why we use the FG model instead of standard or canonical models, which are defined without the variable *σ* [30–34]. The problem is whether or not the results of the canonical 2D and 3D models are identical to those of the FG models. The answer to this question is that the FG model is more suitable than the canonical model. In fact, almost the same results are obtained for some limited range of *κ*; however the stress-strain diagram obtained from the canonical 2D model for a certain finite value of *κ* is not J-shaped, as reported in Ref. [28]. Another reason is that only the FG model reproduces the diagrams that are convex upwards for the large-strain region. These are the reasons for why we use the FG modeling technique to analyze the experimental J-shaped diagrams. In addition to these features of FG modeling, it is important to note that the mechanical strength of real membranes, e.g., the surface tension, becomes dependent on the position and direction on the surface [3, 5]. This property is the origin of why the FG model is considered more suitable for actual biological membranes than the standard surface models [35].

## II. MODELS AND MONTE CARLO SIMULATIONS

We use cylindrical surfaces that are suitable for the calculation of the surface tension or the stress. The 2D FG model is defined on a cylindrical lattice like the one in Fig. 2(a). The size of the lattice is given by (*N*, *N*_*B*_, *N*_*T*_) = (2511, 7371, 4860), where *N*, *N*_*B*_, and *N*_*T*_ are the total number of vertices, the total number of bonds, and the total number of triangles, respectively. The Euler number *χ* is the same as that of the torus, and hence, *χ* = *N - N*_*B*_ + *N*_*T*_ = 0. The lattice in Fig. 2(b) is constructed from tetrahedrons for the 3D FG model [36], and the surfaces inside and outside the 3D lattice are 2D cylinders that are exactly same as the one in Fig. 2(a) [37]. This 3D lattice is thin, and hence, all vertices are on the surface; there are no vertices inside the 3D structure. The lattice size is given by (*N*, *N*_*B*_, *N*_*T*_, *N*_tet_) = (5022, 24624, 34182, 14580), where the first three symbols are the same as those for 2D lattice and *N*_tet_ is the total number of tetrahedrons. This 3D lattice is topologically identical to the torus and has zero Euler number *χ* = *N – N*_*B*_+*N*_*T*_ – *N*_tet_ = 0. The height *H* is fixed during the simulations for the calculation of the tensile force (Fig. 3(a)). The diameter *D* of the upper and lower boundaries is also fixed to *D*_0_, and this boundary condition protects the cylinder from collapsing for small bending rigidity in the simulations. It should be emphasized that this constraint for *D* on the boundaries is close to the experimental setup for the measurement of tensile force.

**FIG. 2.**
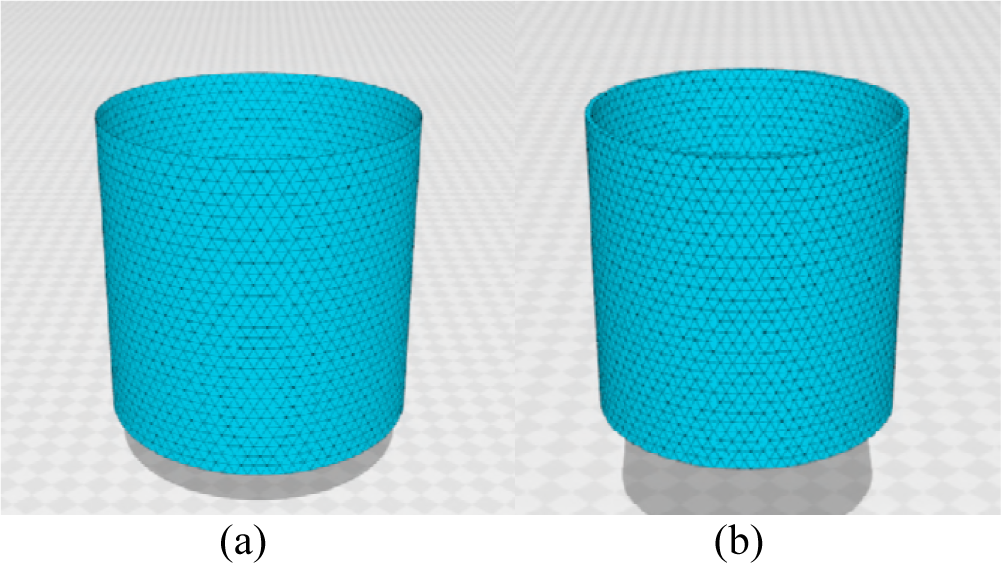
Cylindrical lattices for (a) 2D and (b) 3D FG models. The lattices are composed of (a) triangles and (b) tetrahedrons. The total number of vertices are (a) *N* = 2511 and (b) *N* = 5022.

**FIG. 3.**
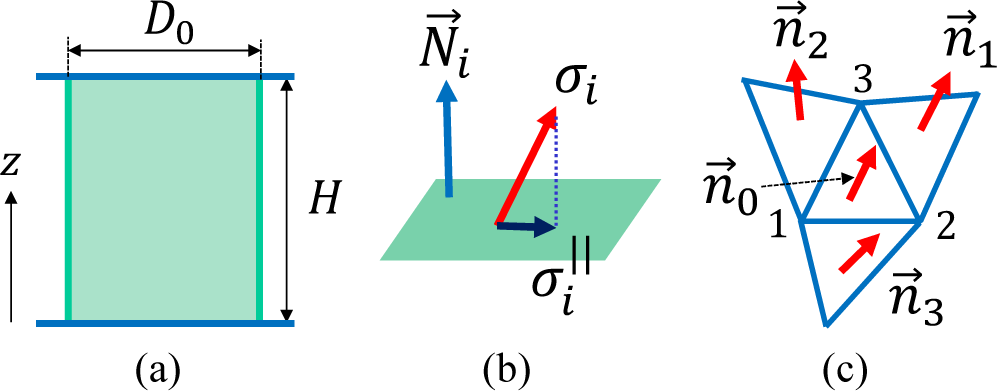
(a) The height *H* and diameter *D*_0_ of a cylinder, the unit normal vector **N**_*i*_ of the tangential plane at the vertex *i*, the tangential component 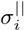 of the variable *σ*_*i*_, and the unit normal vectors **n**_*i*_ of the triangles *i*(= 0, 1, 2, 3) used in *S*_2_ of Eq. (4).

### A. 2D model

In this subsection, we describe the 2D FG model. Although this 2D model is the same as the one in Ref. [28], we summarize the Hamiltonian here in a self-contained manner. The Hamiltonian is given by a linear combination of four terms such that

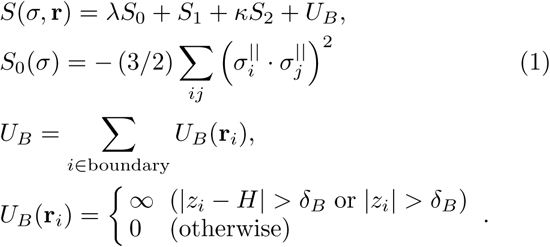

The variable **r**(*∈* **R**^3^) is the vertex position and represents the position of a polymer, such as collagen fibers. The direction of the polymer is represented by *σ*(*∈ S*^2^), which has a non-polar interaction of the Lebwohl-Lasher type [38] described by *S*_0_; hence, *σ* is identified with *-σ*. We should note that the edges of the triangles do not always represent linear polymers or polymer networks [39]. The triangles are simply introduced for the discretization of 2D materials [40–45]. Indeed, the triangle edges play a role as local coordinate axes for the discretization of the Hamiltonian. The variable 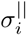 in *S*_0_ is defined by

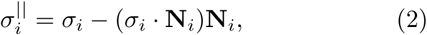

which is a component of *σ*_*i*_ parallel to the tangential plane at the vertex *i* (Fig. 3(b)). This tangential plane is determined by its unit normal vector **N**_*i*_ (Fig. 3(b)), which is defined such that

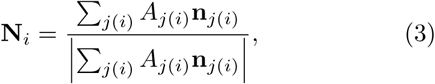

where *A*_*j*(*i*)_ and **n**_*j*(*i*)_ denote the area and the unit normal vector of the triangle *j*(*i*) sharing the vertex *i*, respectively (Fig. 3(c)).

The Gaussian bond potential *S*_1_ and the bending energy *S*_2_ are given by

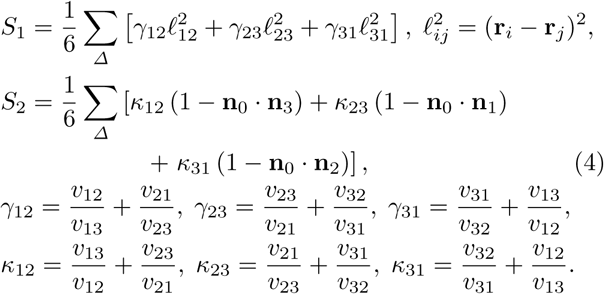

In these expressions, *ℓ*_*ij*_ is the length of bond *ij* connecting the vertices *i* and *j*, and **n**_*i*_ is a unit normal vector of the triangle *i*. The coefficient *γ*_*ij*_ in *S*_1_ is defined by using *v*_*ij*_, which is given by

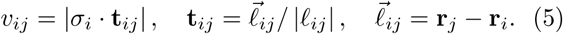

(see Ref. [36] for more detailed information on the discretization of *S*_1_ and *S*_2_). The quantities *γ*_*ij*_(= *γ*_*ji*_) and *κ*_*ij*_(= *κ*_*ji*_) are considered the positionand directiondependent surface tension and bending rigidity, respectively.

The potential *U*_*B*_ allows the boundary vertices to move vertically in the *z*-direction within a small range ± *δ*_*B*_, which is fixed to the mean bond length. This constraint does not influence the results in the limit of *N µ ∞* because *ξ /H* is negligible in this limit

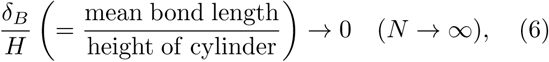

because the mean bond length is independent of *N*, while *H* is proportional to *N*. The reason for this constraint *U*_*B*_ is assumed to be avoiding a strong and non-physical force, which is suspected to appear when *σ*_*i*_ aligns with the *z*direction on the boundary. If *σ*_*i*_ on the boundary aligns with the *z*-direction without *U*_*B*_, the corresponding *v*_*ij*_ becomes *v*_*ij*_ *→* 0 because of the definition of *v*_*ij*_ in Eq. (5). Therefore, the corresponding *γ*_*jk*_ *→ ∞* becomes *γ*_*jk*_, and hence, *S*_1_ → *∞*. In this situation, the variable *σ*_*i*_ never aligns with the *z*-direction, and therefore, *U*_*B*_ is necessary for the well-definedness of the model.

The partition function is given by

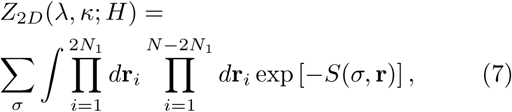

where *H* is the height of the cylinder and is fixed during the simulation (Fig. 3(a)).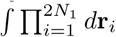 denotes the 1Di integrations of the vertices on the boundaries, where *N*_1_ is the total number of vertices on the upper and lower boundaries. The 2*N*_1_ vertices are allowed to move along the circles of radius *D*_0_, and hence, the corresponding integration effectively becomes one-dimensional. The total number of remaining vertices is *N* – 2*N*_1_, and the positions of these vertices are integrated out by the 3D integrations represented by 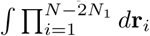.

### B. 3D model

The 3D model Hamiltonian is defined on the 3D lattice discretized by the tetrahedrons shown in Fig. 4(a) [37]. Although the Hamiltonian is almost the same as that in Ref. [36], we briefly describe it in the outline below. The model is defined without the self-avoiding potential for the ”surfaces” (not for the inside of the structure), and this is the only difference between the models in this paper and in Ref. [36]. The surface self-avoiding potential is a non-local potential and is time consuming for simulations. We expect that the results are not strongly influenced by whether this self-avoiding interaction is included or not because the upper and lower boundaries are fixed and the surface always remains relatively smooth. This is also expected for the 2D model, which has no self-avoiding potential.

**FIG. 4.**
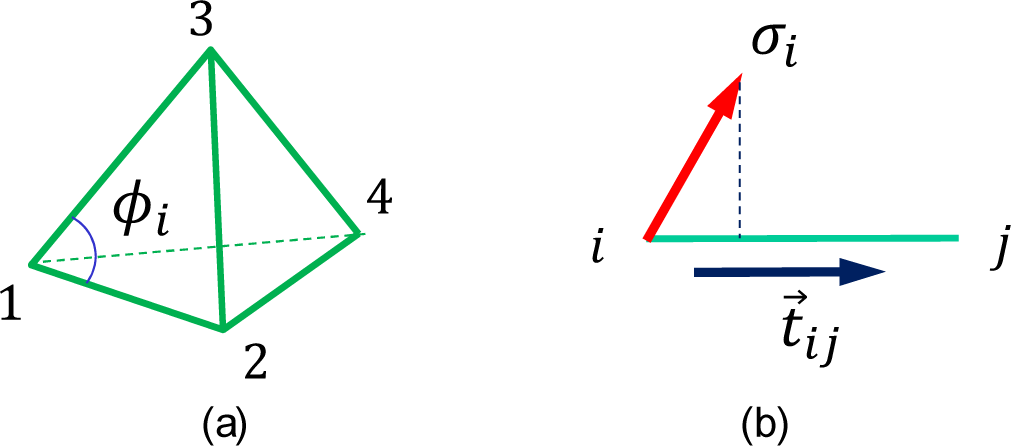
(a) A tetrahedron with vertices (1,2,3,4) and an internal angle *ϕi* of a triangle and (b) the variable *σi* and the unit tangential vector **t**_*ij*_ of the bond connecting the vertices *i* and *j*. The *σi* and **t**_*ij*_ are used to define *v*_*ij*_ in Eq. (5).

The Hamiltonian *S*(**r**, *σ*) is defined by a linear combination of five different terms

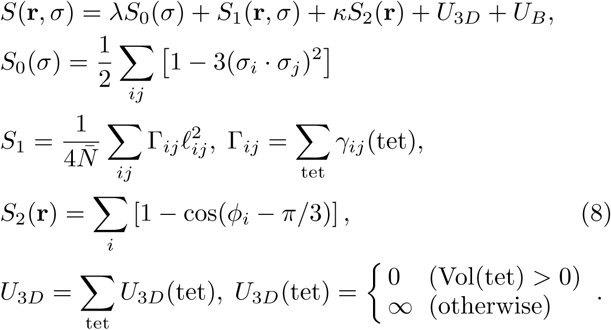

The variable **r**(*∈* **R**^3^) is the vertex position of a tetrahedron, and *σ*(*∈ S*^2^) denotes the directional degrees of freedom of polymers exactly the same as in the 2D model. Each term shares the same role with the corresponding term in the 2D model. The definition of the Lebwohl-Lasher potential *S*_0_ is slightly different from that of *S*_0_ in Eq. (1), but the role of this term in the 3D model is identical to that of *S*_0_ in the 2D model. The definition of *S*_1_ is also slightly different from the 2D case; however, the continuous description of *S*_1_ is the same (see Ref. [36]). In the coefficient of *S*_1_,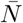 is defined by

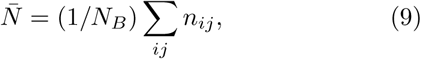

where *n*_*ij*_ is the total number of tetrahedrons sharing the bond *ij* and *N*_*B*_(=∑_*ij*_ 1) is the total number of bonds. The *γ*_*ij*_(tet) in *S*_1_ is given by

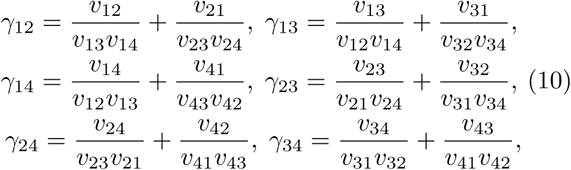

where the numbers 1, 2, 3, 4 denote the vertices of the tetrahedron in Fig. 4(a). In these expressions for *γ*_*ij*_, the symbol *v*_*ij*_ is defined by the same expression as in Eq. (5) using *σ*_*i*_ and the unit tangential vector **t**_*ij*_ along the tetrahedron edge *ij* (Fig. 4(b)).

The term *S*_2_ in Eq. (8) is different from that of the 2D model in Eq. (1); however, the role of *S*_2_ in Eq. (8), i.e., to keep the tetrahedron shape almost regular for positive *κ* values, is the same as that in the 2D model. The symbol *ϕ*_*i*_ is the internal angle of triangles (Fig. 4(a)). The role of the potential *U*_3*D*_ is to protect the tetrahedron volume from being negative. This potential *U*_3*D*_ introduces a repulsive interaction between the vertices so that the tetrahedron is hardly collapsed, and hence, this *U*_3*D*_ shares the same role with *S*_2_ in part. For this reason, the tetrahedron hardly deforms for positive *κ* values, and therefore, we assume small negative *κ* values in the simulations to make the J-shaped diagrams have a large strain. The potential *U*_*B*_ is exactly same as in Eq. (1) for the 2D model, and for this reason, its definition is not written in Eq. (8).

The partition function *Z*_3*D*_ can also be defined for the 3D model; however, its description is exactly same as *Z*_2*D*_ in Eq.(7) except the actual number *N*_1_ for the boundary vertices; to avoid duplication, it is not written here.

### C. Formula for stress calculation

In both the 2D and 3D models, the stress in the stress strain diagram is calculated from the principle of scale invariance of the partition function *dZ/dα*|_*α*=1_ = 0 (for simplicity, the subscript 2*D* in *Z*_2*D*_ is not written henceforth) [46]. This invariance simply originates from the fact that the integrations in *Z* are independent of its expression for **r**, the position of the material.

If we change **r** to *α***r** with a positive number *α*, we have the scaled partition function such that

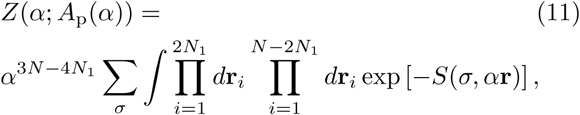

where *A*_p_ is the projected area of the surface and *N*_1_ is the total number of boundary vertices, as mentioned in the previous subsection. The expression *Z*(*α*; *A*_p_(*α*)) implies that *Z* depends both explicitly and implicitly on *α*. In the Hamiltonian *S*(*σ, α***r**), the only term that depends on *α* is *S*_1_: *S*_1_(*α*) = *α*^2^*S*_1_. We should note that the coefficient *α*^3^*N-*^4^*N*^1^ in the right hand side of Eq. (11) comes from the 3D and 1D integrations such that *α*^3*N -*4*N*_1_^ = *α*^3(*N -*2*N*_1_)^*α*^2*N*_1_^.

From the abovementioned scale invariance of *Z*, we have *d* log *Z/dα|*_*α*=1_ = 0 and

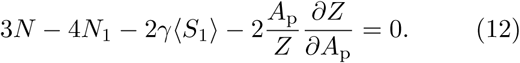

For the last term on the left hand side, we assume that *A*_p_(*α*)= *α*^-2^*A*_p_ for the dependence of *A*_p_(*α*) on *α* because the projected area *A*_p_ is kept fixed under the scale change **r** → *α***r** for the evaluation of the tensile stress *τ* (Fig. 5(a)). Moreover, to evaluate *∂Z/∂A*_p_ on the left hand side of Eq. (12), we naturally assume that the surface is sufficiently expanded. Under this condition, the free energy *F* of the surface is given by

**FIG. 5.**
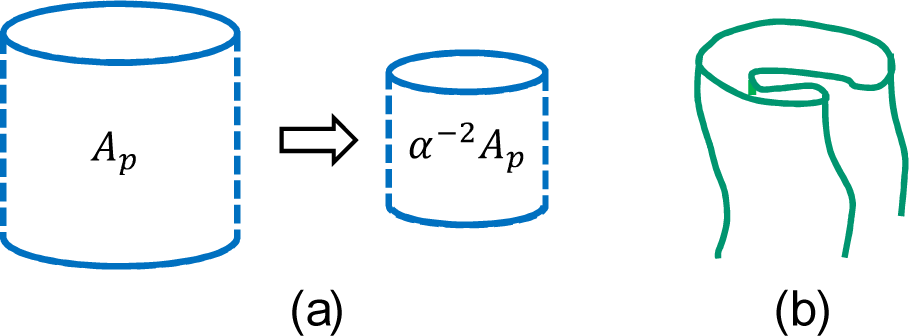
(a) An illustration of the change in the projected area *α*^-2^*A*_p_, which restores the original *A*_p_ and remains unchanged under the scale change **r** *→ α***r** in the partition function, and (b) a possible folding of the surface (which is magnified; this type of folding is always suppressed).

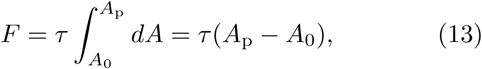

where *A*_0_ is the area of the surface corresponding to the zero-tensile force [46]. Thus, we have the partition function *Z* = exp(*-F*). Inserting this *Z* into Eq. (12), we have

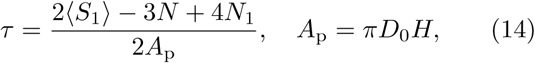

where *D*_0_ is the diameter of the boundary. We should note that 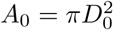, where the initial height is given by *H* = *D*_0_. This surface tension *τ* in Eq. (14) is called the frame tension because *τ* depends only on the area of the frame on which the surface spans. We should note that the formula for *τ* of the 3D model is the same as Eq.(14) for the 2D model. This is because the thickness of the 3D cylinder the for 3D model shown in Fig. 2(b) is sufficiently thin that this 3D cylinder is regarded as a 2D surface.

The reason why *D*_0_ = *H* is assumed in the configurations for *τ* = 0 is because the lattice is constructed under the condition *D*_0_ = *H* with a regular triangle (see Fig. 2). The edge length is expected to be uniform and independent of the direction in the initial undeformed configuration if *D*_0_ = *H* is satisfied, at least for *λ* → 0. Therefore, we have no reason to fix *D*_0_, for example, *D*_0_ ≠ *H*, although a non-zero *λ* is assumed in both the 2D and 3D simulations.

To compare the simulation result *τ* in Eq. (14) with the experimental data, we have to change simulation units to physical units. For this purpose, we explicitly use *k*_*B*_*T* and the lattice spacing *a* [28, 47], which are suppressed by *k*_*B*_*T* = 1 and *a* = 1 in the expression for *τ* in Eq. (14). All quantities that have units of length are multiplied by *a*, and the Boltzmann factor exp(–*F*) is replaced by exp(–*βF*), where *β* = 1*/k*_*B*_*T*. It should also be noted that *τ* is the surface tension and has units of [N/m], whereas the experimentally measured stress has units of [N*/*m^2^]. Because of this difference in the units, the simulation data *τ* should be divided by *a* when comparing them to the experimental data. Thus, we have the expression *τ*_sim_ for the simulation data with units of [N*/*m^2^]:

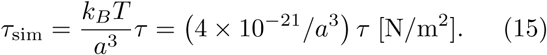

In this expression, *a* is varied to modify the simulation data *τ*, and the modified *τ*_sim_ can be compared to the experimentally observed stress *τ*_exp_.

The Young’s modulus *E* can also be determined from the linear region of the simulation data *τ* by dividing *τ* by the strain. The obtained *E* is modified by the same factor in *τ*_sim_ such that

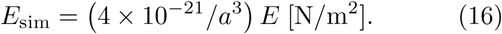

This *E*_sim_ is directly compared to the experimental Young’s modulus *E*_exp_, which is called the stiffness and determined from the linear region of *τ*_exp_ [3, 6–8]. The value of *a* in this *E*_sim_ is the same as that of *a* in *τ*_sim_. Therefore, *E*_exp_ is not independent of *τ*_exp_. For this reason, we compare only *τ*_exp_ with *τ*_sim_ in this paper.

We should note that among the parameters used in the simulation, not all of them are always comparable to physical quantities. The only quantities that can be compared to the experimental ones are *τ*_sim_ and *E*_sim_. In fact, we assume that *κ* is negative in the 3D FG model. The reason for the negative *κ* is that the tetrahedrons hardly deform for positive *κ* values, i.e., where the obtained diagram has no toe region, as mentioned above.

### D. Monte Carlo technique

The standard Metropolis MC technique is used to update the variables **r** and *σ* [48, 51]. The variable *σ* is updated by using three different uniform random numbers, and the new variable *σ*^*′*^ is defined independently of the old *σ*. The variable **r** is updated such that **r** → **r**^*′*^ = **r**+*δ***r** with a small random vector *δ***r**. This vector *δ***r** is randomly generated in a sphere of radius *d*_0_, which is fixed for an approximately 50% acceptance rate.

On the upper and lower boundaries, the new position **r**^*′*^ is constrained such that the diameter of the boundaries remains constant at *D*_0_, as mentioned above. For the 3D model, two different diameters, 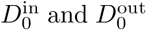, are assumed: one for the inner cylinder and the other for the outer one. The difference is given by 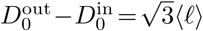, where ⟨*ℓ*⟩ is the mean bond length of the initial configuration for the simulation. In the discussions below, we use only *D*_0_ for simplicity. The constraint on the diameter allows the vertices to move only along the circles of diameter *D*_0_. Due to this free movement of vertices along the circle, it is possible that the surface will become folded for a range of relatively small *κ* values when the height *H* is sufficiently small and close to *D*_0_ (Fig. 5(b)). However, as will be seen in the snapshots of the surfaces, no folding is expected.

Another constraint is imposed on the boundary vertices by *U*_*B*_ in Eq. (1). Under this *U*_*B*_, the vertex positions can move in the vertical direction (⇔ *z*-direction) within the small range *δ*_*B*_. This *δ*_*B*_ is fixed to the mean bond length, as mentioned in the previous subsection.

The lattice size for 2D simulations is (*N, N*_*B*_, *N*_*P*_) = (10584, 31416, 20832), and the size for 3D simulations is (*N, N*_*B*_, *N*_*T*_, *N*_tet_) = (9761, 48124, 66965, 28602). For this 3D lattice, which is constructed using the same technique as the lattice in Fig. 2(b), there are no vertices inside the structure, and all the vertices are on the surface.

## III. SIMULATION RESULTS

### A. Comparison with experimental data

We show in Fig. 6 the experimental stress-strain data of snake’s skin reported in Ref. [3]. The skin of snakes, which is composed of collagen and elastin fibers, has a relatively large deformation, as expected from their typical body elongation and bending. The units of the stress *τ*_exp_ are [MPa], and the snake skin is relatively strong. The toe region ranges from 50% to 125% depending on the body position from which the skin is sampled. The sampling positions of the data (Exp) in Fig. 6(a) and Fig. 6(b) are 40% and 60% distant from the snake’s snout, where 100% corresponds to the length between the snout and the vent. The toe length of the plotted data in Figs. 6(a),(b) is relatively large and almost equal to 100% and 125% in units of strain. These curves are typical examples of diagrams with an S-heel, as mentioned in the Introduction, and they are composed of two different linear lines. The parameters used for the simulations are summarized below in Table I.

**FIG. 6.**
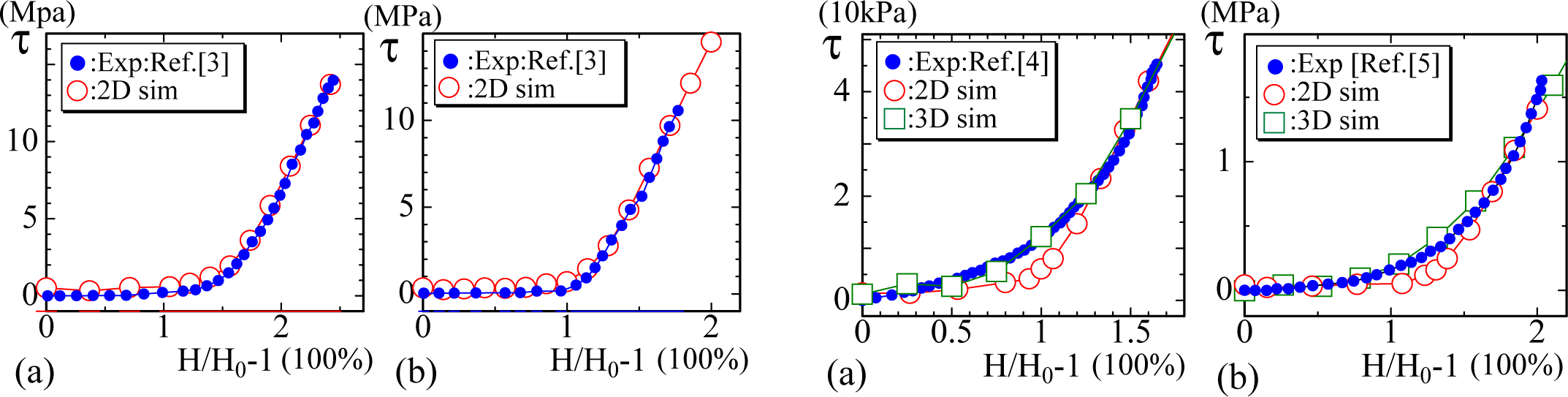
Comparison of experimental data *τ*_exp_ (Exp) with the simulation data *τ*_sim_ (sim) for snake skin with a toe region of 100% and (b) 125% in units of strain [3]. These experimental data are examples of diagrams with an S-heel, and the 2D simulation results are used for comparison with these experimental data.

The stress *τ*_sim_ in the 2D model in Fig. 6(a) is calculated by Eq. (15) using the simulation data *τ*, and *τ*_sim_ is found to be almost identical to the experimental data *τ*_exp_. The parameter *λ* in Eq. (1) is fixed to *λ* = 1, and the bending rigidity *κ* is varied for the simulations of the 2D model in this paper. The assumed bending rigidities are *κ* = 0.6 and *κ* = 0.55 for the 2D simulations in Figs. 6(a) and 6(b), respectively. The lattice spacing *a* in Eq. (15) used for the fitting is *a* = 0.83 × 10^−9^ in Fig. 6(a) and *a* = 0.85 × 10^−9^ in Fig. 6(b), both of which are larger than the Van der Waals radius (∼1 × 10^−10^[m]), i.e., the typical size of atoms. This is the reason why we call the FG model a coarse-grained model.

The second and third sets of experimental data are of soft biological tissues such as diaphragm and arteries, to which the 2D model data are not always well fitted. The data plotted in Fig. 7(a) are those obtained in the study of the mechanical diaphragm in a model of muscular dystrophy [4]. The data in Fig. 7(b) are the diagram measured along the circumferential axis of arteries [5]. In Ref. [5], Arroyave et al. investigated a methodology for mechanical characterization of soft biological tissues. In that work, biaxial tensile testing was performed, and methodologies for testing and data processing were proposed. Sections of three groups of arteries, such as the abdominal aorta, thoracic aorta and left subclavian artery, were selected from pigs in their study. They also analyzed the experimental data by using continuous mechanical models (Fung’s Model and Holzapfel’s Model). It was found that abdominal aortas can support higher stress before rupture due to the presence of collagen in the samples. It is also possible to understand that high percentages of elastin are the reason for the large strain in all groups.

**FIG. 7.**
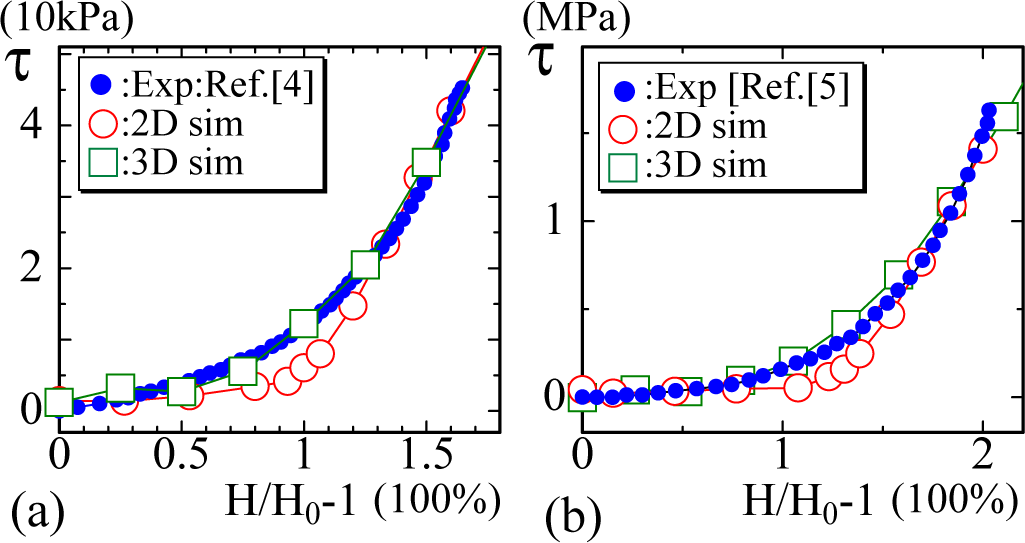
The experimental diagrams of soft biological tissues such as (a) skin and (b) arteries reported in [4, 5]. These experimental data are examples of diagrams with an L-heel, and the 2D simulation data slightly deviate from the experimental data in both (a) and (b).

The curvature of the heel region in both experimental curves plotted in Figs. 7(a),(b) is relatively small compared to that in the experimental data shown in Figs. 6(a),(b). For this reason, the fitting of the simulation data of the 2D FG model is not good for these data, while the 3D simulation data are well fitted except in the failure region.

The next experimental data reported in Ref. [6] are of the periodontal ligament of the molars of rats at (a) 6 months and (b) 12 months of age (Figs. 8(a),(b)). In the case of this material, the strain is over 250% and is larger than that in the previous examples shown in Figs. 6 and 7. Moreover, for the large-strain region, the curve of the experimental data starts to bend and becomes convex upwards. The simulation data of 2D model are well fitted even for the convex part (except the failure region), although the 3D simulation data start to deviate from the experimental data in the convex region in Fig. 8(a). We should emphasize that only the 2D FG model produces results that are in good agreement with the experimental data. In fact, the results of the canonical model are always linear for the large-strain region, and they cannot be fit to the experimental curve with the convex part. This is a non-trivial difference between the FG and canonical models, as is the fact that the diagram of the canonical model becomes linear at a certain value of *κ*, while the diagram of the FG model is always J-shaped independent of *κ* [28].

**FIG. 8.**
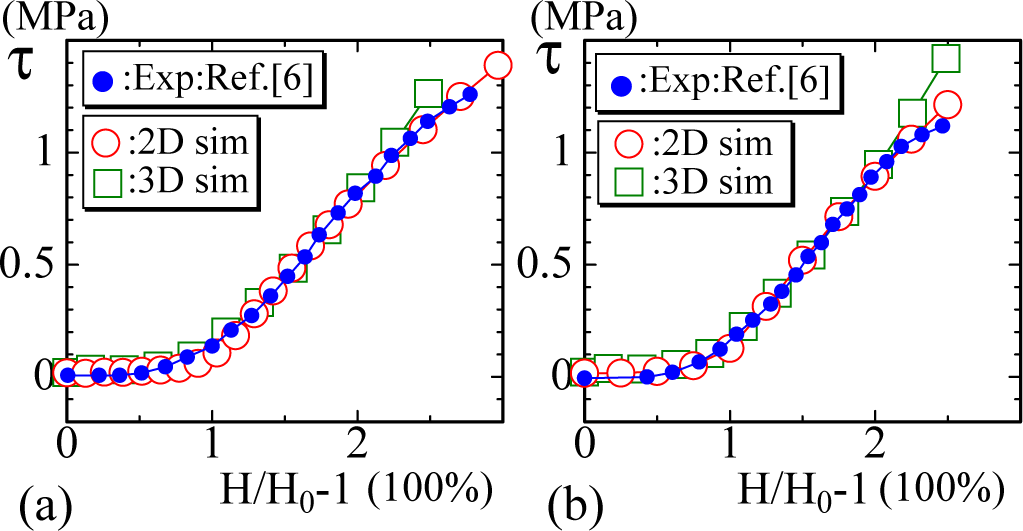
The experimental data are of the periodontal ligament of the molars of rats at (a) 6 months and (b) 12 months of age [6]. The strain of the toe region is approximately 50%, which is relatively short. For the large-strain region, the experimental data are slightly convex upwards, which is well fitted by the 2D simulation data except in the terminal failure region.

The final examples of experimental data shown in Figs. 9(a),(b) are of the passive tension vs. titin strain of rat muscles, namely, skeletal and cardiac muscles [7]. Granzier et al. studied the mechanical properties of cardiac muscle by investigating passive tension and stiffness in a stretch-and-release process. They estimated the contribution of collagen, titin, microtubules, and intermediate filaments to the tension and the stiffness by gluing experiments after chemical treatments. Because irregular isoforms of certain proteins might cause heart diseases such as dilated cardiomyopathy, as reported earlier [49, 50], it is important to understand why cytoskeletons play significant roles in passive tension and how they function. In their study, it was found that passive tension of cardiac cells increases slowly at the shorter end of the working range (1.9 to 2.2 *µ*m) in the heart but increases more steeply at longer lengths up to 4.5 *µ*m. They concluded that titin and collagen are the best two candidates for explaining the experimental results over a wide range of the lengths. On the other hand, microtubules and intermediate filaments do not develop passive tension significantly in the working range. In addition, as opposed to passive tension in skeletal muscles, cardiac passive tension does not show a plateau at long lengths around 4 *µ*m. However, they argued that this plateau can be observed by excluding the effect of intermediate filaments.

**FIG. 9.**
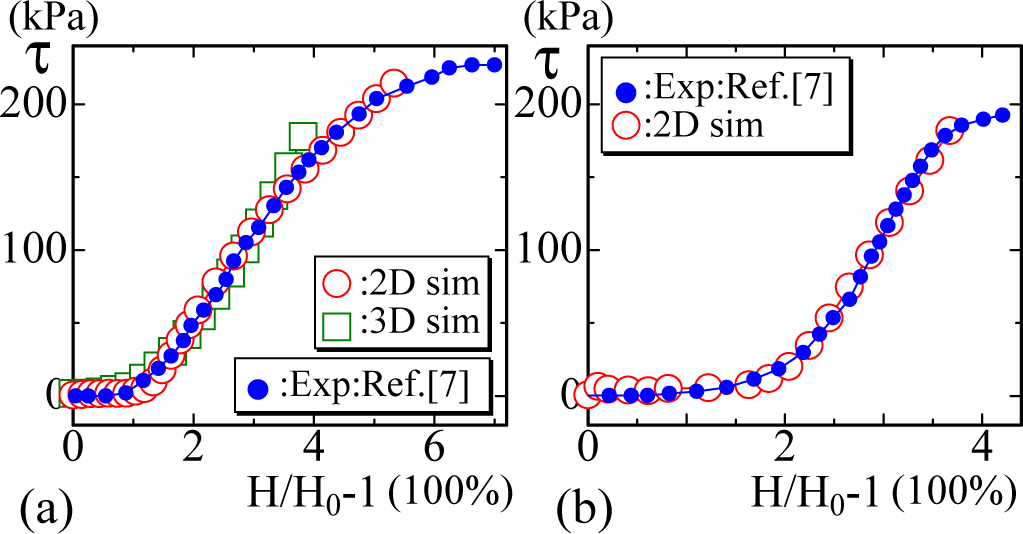
The stress vs. titin strain of animal’s (a) cardiac and (b) skeletal muscles, where the range of the toe region extends to 100% and 150% [7] and the curves have a convex part in the large-strain region. The strains are very large (up to 700%) compared to those of the other materials studied in this paper.

These materials are very soft and flexible, and the titin strain is very large and ranges from 400% to 700%. The experimental curves are convex upward in the large-strain region, and the convex shape is more clear than that in the data shown in Figs. 8(a),(b). In addition to the previous data with the convex part, the results of the 2D FG model better fit the experimental data.

The parameters assumed in the simulations are summarized in Table I. All values of *a* are sufficiently larger than the Van der Waals radius. These are the microscopic parameters and do not always correspond to actual physical quantities, as mentioned in Section II C.

**TABLE I.** The parameters assumed for the 2D and 3D models. The units of the parameters are *λ*[*β*], *κ*[*β*], *D*_0_[*a*] and *a*[m], where *β* = 1*/k*_*B*_ *T*.

| | Model | $\lambda$ | $\kappa$ | $D_0$ | $a$ |
| --- | --- | --- | --- | --- | --- |
| Fig.6(a) | 2D | 1 | 0.6 | 14 | $0.83 \times 10^{-9}$ |
| Fig.6(b) | 2D | 1 | 0.55 | 11.7 | $0.85 \times 10^{-9}$ |
| Fig.7(a) | 2D | 1 | 0.6 | 15 | $0.52 \times 10^{-8}$ |
| | 3D | 0.4 | -0.05 | 8 | $0.75 \times 10^{-8}$ |
| Fig.7(b) | 2D | 1 | 0.6 | 13 | $0.17 \times 10^{-8}$ |
| | 3D | 0.45 | -0.05 | 7.6 | $0.25 \times 10^{-8}$ |
| Fig.8(a) | 2D | 1 | 0.6 | 15.5 | $0.23 \times 10^{-8}$ |
| | 3D | 0.4 | -0.05 | 8.6 | $0.34 \times 10^{-8}$ |
| Fig.8(b) | 2D | 1 | 0.6 | 16 | $0.23 \times 10^{-8}$ |
| | 3D | 0.4 | -0.05 | 8.6 | $0.33 \times 10^{-8}$ |
| Fig.9(a) | 2D | 1 | 0.6 | 13.5 | $0.52 \times 10^{-8}$ |
| | 3D | 0.35 | -0.1 | 8.3 | $0.83 \times 10^{-8}$ |
| Fig.9(b) | 2D | 1 | 0.5 | 9.8 | $0.44 \times 10^{-8}$ |

### B. Behavior of the variable *σ* and snapshots

To observe how the variable *σ* aligns, we calculate the order parameter *M* of *σ* by

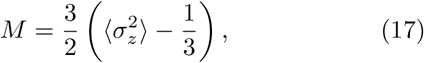

which represents the alignment of *σ* along the *z* axis [38]. We also calculate the eigenvalues of the tensor order parameter

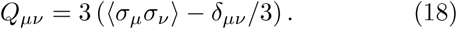

The largest eigenvalue ∑_1_ of *Q*_*µv*_ and *M* corresponding to several simulation results is plotted in Figs. 10(a),(b). For the small-strain region, S_1_ and *M* slightly deviate from each other; however, they are exactly the same for the large-strain region. This implies that *σ* aligns in the *z*-direction to which the tensile force is applied.

**FIG. 10.**
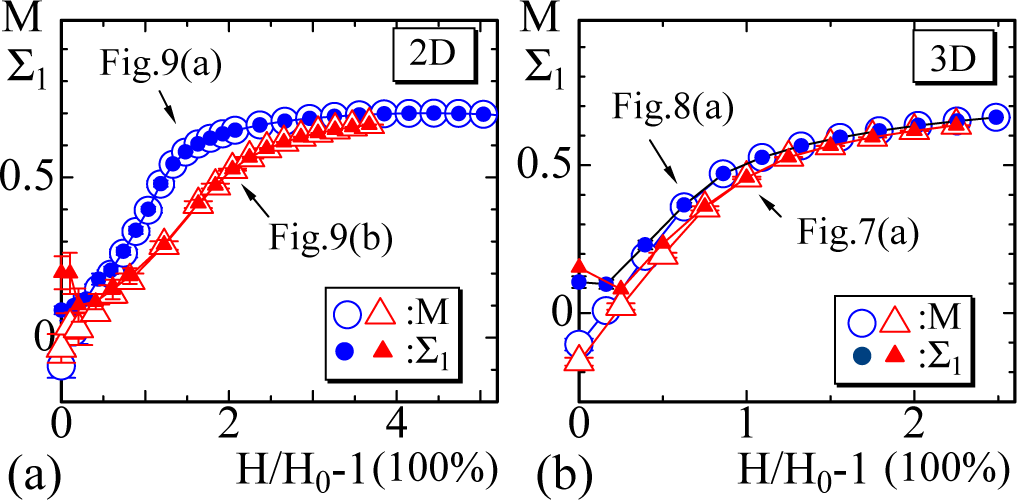
The open symbols in (a) 2D and (b) 3D simulations represent the order parameter *M* defined by Eq. (17), and the solid symbols represent the largest eigenvalue S_1_ of the tensor order parameter defined by Eq. (18).

Snapshots of the surfaces are shown in Figs. 11(a)-(h), where the scales of the figures are different from each other because the height difference is very large. We find that the variable *σ* locally aligns at least in the configurations for *H* = *D*_0_ in Figs. 11(a) and 11(e). This implies that *σ* changes in a similar manner to the directional degree of freedom of polymers, which undergo a transition from a locally ordered polydomain phase to a globally ordered monodomain phase. We should note that the locally ordered configuration in the small-strain region is not always random but almost ordered. In fact, a configuration of *σ* with a circular direction, for example, has no contribution to *M* because the surface is cylindrical. However, it is expected that this almost-ordered phase is consistent with the fact that linear objects such as collagen fibers in biological tissues are almost aligned even for the released and zero-strain configurations.

**FIG. 11.**
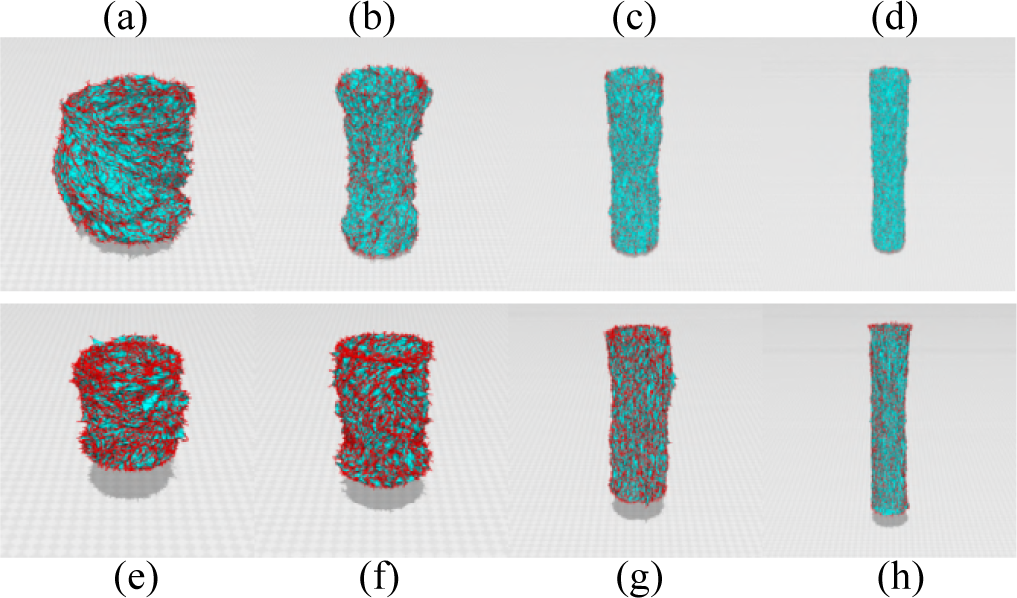
Snapshots of cylindrical surfaces of the 2D model ((a)-(d)) and 3D model ((e)-(h)) corresponding to the simulation data in Fig. 9(a). The heights of surfaces are (a) *H*(= *D*_0_) = 14, (b) *H* = 32, (c) *H* = 52, and (d) *H* = 78 for the 2D model and (e) *H*(= *D*_0_) = 8.3, (f) *H* = 12, (g) *H* = 24, (h) *H* = 40 for the 3D model. The scales of the figures are different from each other. The (red) burs on the surface represent the variable *σ*.

## IV. SUMMARY AND CONCLUSION

We studied the large-strain J-shaped diagrams of biological membranes such as animal skin, muscles and arteries by 2D and 3D Finsler geometry (FG) models. These materials are very soft, and the zero-stress region of the diagrams ranges from approximately 50% to 150%. Because of this zero-stress region, the diagram is called J-shaped. The J-shaped diagram is roughly composed of two linear lines except the failure region: one is the toe region, and the other is a linear region. The region where these two lines are smoothly connected is called the heel. Based on the word ”heel”, we can divide the experimental large-toe diagrams into two groups: diagrams with a small heel (S-heel) and diagrams with a large heel (Lheel). The diagrams with an S-heel are consistent with the results of the 2D model, while the diagrams with an L-heel are well fitted by the results of the 3D model. Moreover, the experimental diagrams with a convex part in the large-strain region can be fitted by the 2D FG simulation data. These observations show that the FG modeling technique is suitable for analyzing J-shaped diagrams of biological membranes.

The FG model is constructed by extending the linear chain to a 2D surface or a 3D body. In the 2D and 3D models, a Finsler metric is assumed in the Gaussian bond potential, and this Finsler metric plays a role in introducing the interaction between the variables *σ* and **r**, which correspond to the polymer direction and position, respectively. As a result of this modeling, the mechanical strength, such as the surface tension, effectively depends not only on the position but also on the direction inside the material. This is the main advantage of the FG model over its canonical counterpart. We should note that this FG modeling is only possible for 2D and 3D models because 1D space is a line with no non-trivial metric function.

The fact that the FG model was successfully applied to large-strain J-shaped diagrams indicates that highly nonlinear large-strain diagrams of polymers, include those with rubber elasticity, can also be targeted [39, 52]. This will be a biology-inspired challenge in understanding new materials and their possible mechanical properties.

## V. AUTHOR CONTRIBUTIONS

K.M. and S.G. studied and analyzed the existing experimental data and wrote the presentation section of the manuscript. H.K. performed the simulations and wrote the other part of the manuscript.

## ACKNOWLEDGMENTS

The authors acknowledge Y. Takano for computer analyses. This work is supported in part by JSPS KAKENHI Grant No. 17K05149.

